# Discovery of Novel Trypanothione Reductase Inhibitors through Pharmacophore Modeling, virtual Screening and Molecular Dynamics Simulations. New insights for Human African Trypanosomiasis

**DOI:** 10.1101/2025.04.18.649577

**Authors:** Elcanah Mauta Evans, Oluwaseun Oluwatosin Taofeek, Theresa Manful Gwira, Collins Misita Morang’a, Thommas Mutemi Musyoka, Steven Ger Nyanjom

## Abstract

Human African Trypanosomiasis (HAT), caused by *Trypanosome brucei*, remains a critical health concern, with treatments such as Melarsoprol hampered by toxicity and resistance. To address this, we aimed to identify inhibitors targeting Trypanothione Reductase (*T.b* TR), a key enzyme in the parasite’s survival. A pharmacophore model was derived from Melarsoprol and validated using Receiver Operating Characteristic (ROC) curves and Enrichment Factor (EF). Virtual screening of compound libraries identified two promising candidates, VS-1 and VS-2, based on superior docking scores. Subsequent molecular dynamics simulations confirmed the stability of the ligand-enzyme complexes, while binding free energy calculations revealed strong binding affinities for VS-1 and VS-2. Per-residue decomposition pointed critical interactions with active site residues, including MET 260, ASN 130, and HIS 128, contributing to the compounds’ stability and activity. These findings suggest that VS-1 and VS-2 are potential inhibitors of *T.b* TR, with better efficacy than Melarsoprol. However, further *in vitro* studies, including IC_50_ determination, cell cycle analysis, and morphological assays, are essential to confirm the therapeutic potential of these compounds for the treatment of HAT.

## Introduction

Human Animal Trypanosomiasis is a Tropical neglected disease of global concern, especially in the sub-Saharan Africa [1]. The disease takes two forms depending on the *Trypanosome brucei* subspecies responsible for the transmission. These subspecies include *Trypanosome brucei gambiense,* which is responsible for 92% of cases and is found in 24 African countries, particularly in the west and central Africa and *Trypanosome brucei rhodesiense* which is found in 13 sub-Saharan countries, specifically in the Eastern and southern Africa [2]. During haemolymphatic stage, pentamidine or suramin are administered since the parasites are distributed in the blood stream, organs, and adipose tissues. However, in later stages (meningo-encephalitic) pentamidine or suramin cannot be used as the parasites have entered the central nervous system. This is because these drugs cannot cross the blood brain barrier to reach the parasites [3]. Melarsoprol is commonly used drug in meningo-encephalitic stages of HAT infection. However, the drug has setbacks owing to its toxicity, resistance and administration difficulties [4]. Melarsoprol is an arsenic based drug which is a combination of melarsen oxide and dimercaprol. The drug is metabolized to melarsen oxide which binds to the parasite’s Trypanothione reductase enzyme in a promiscuous nature, hence often reacting with thiol esters of proteins in close proximity, leading to toxicity through oxidative stress [5].

Post treatment reactive Encephalopathic syndrome is a common neurological complication associated with melarsoprol, and it has a high prevalence rate ranging from 2-10% for all patients treated with melarsoprol. Moreover, the case fatality rate of Encephalopathic syndrome is more than 50% [6]. Nifurtimox and eflornithine combination could replace melarsoprol as the combination has much improved efficacy and reduced incidences of drug resistance. However, even though the combination reduces the intervals of drug administrations, the treatment still requires a multiple intravenous administration which is logistically expensive to patients [7]. Recent research shows that an oral monotherapy administration of fexinidazole for a period of 10 days can be used in both early and late stage stages of HAT [7]. However, the drug is not efficient in more severe forms of late stage HAT, therefore, a more aggressive treatment option is still needed.

Structure-based virtual screening is a computational technique used in drug discovery to identify potential drug candidates by screening large libraries of small molecules against the 3D structure of a target protein [8].In this study, we generated a pharmacophore model using melarsoprol, which is the standard drug for late stage HAT. The generated model was then validated using Receiver operating characteristics curve and Enrichment factor before being used for high throughput virtual screening. Multiple ligand molecular docking was carried out using the top scoring ligands from pharmacophore screening. Top two molecules with the least molecular docking affinity scores against *Trypanosome brucei* Trypanothione reductase were selected and validated using molecular dynamics simulations.

## Material and methods

### 1. Pharmacophore querry generation, validation and structural based virtual screening

The standard drug for late stage HAT Melarsoprol was used for pharmacophore querry generation. Pharmit webserver accessible from (https://pharmit.csb.pitt.edu/) was used for pharmacophore modelling. The pharmacophore properties extracted from the drug included a hydrogen Donor (134.7, 117.5, 135.4, r=1), Hydrophobic (134.7, 117.7, 132.5, r=1) and Aromatic property (134.7, 117.7, 132.5, r=1) as shown in figure 1.1. Validation of the modelled pharmacophore querry was done via two steps. First, a library of zinc database was generated using pharmit webserver and screening done using the modelled querry. Search parameters were optimized using molecular weight, number of hydrogen donors/acceptors and tolerance level until the querry retrieved the melarsoprol from the generated zinc database library. This validated the querry since the compound is available in the zinc database. Further validation was done using Receiver operating characteristic curve and enrichment factor. Briefly, 14 active compounds which are known to inhibit *Trypanosome brucei* Trypanothione reductase enzyme which is also inhibited by the standard drug Melarsoprol were picked from literature search. 255 decoys were generated from Database of Useful Decoys webserver available at (https://dude.docking.org/generate). A library of actives and decoys was then generated and uploaded to the pharmit database and the rate at which the generated querry was able to distinguish between actives and decoys during screening estimated using the method previously described by Triballeau and colleagues as shown in figure 1.2 [9]. Briefly, percentage selectivity was defined as the number of selected active compounds per total actives compounds 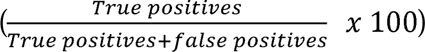. On the other hand, percentage specificity was defined as the total number of discarded inactives per total inactives 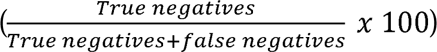. The enrichment factor was computed using (EF = (*number of hit sampled*⁄*hits sample size*)⁄(*Total hits*⁄*Total molecules*).

To carry out high throughput virtual screening the pharmacophore model was further optimized to ensure the number of hydrogen bonds were lower than 4, the molecular weight was set to be within the range of 397 and 399 and only molecules that were able to pass through the enzyme’s active site were considered. Top 20% of the hits were used for validation using the ROC. The databases used for virtual screening included PubChem (103,302,052 molecules), ZINC (13, 127,550 molecules), MCULE (39, 843, 639 molecules), Enamine (4, 117, 328 molecules) and coconut (794, 181 molecules). The results of the top 15 hits from each database screening were saved in an SDF format.

### 2. Molecular Docking, protein-ligand interaction analysis and pharmacokinetic evaluation

The results of the structural based virtual screening across the 5 databases were utilized for multiple ligand molecular docking against *Trypanosome brucei* Trypanothione Reductase enzyme. The enzyme was downloaded from the protein databank using the PDB ID 8JY6.The centroid of the enzyme’s co-crystallized ligand (L1U) was computed using excel and was used as the multiple ligand’s biding site to the enzyme (x=132.27, y=118.06, z=132.18, size x=30, size y=30 and size z =30). Energy minimization was carried out using open Babel’s steepest descent algorithm using MMFF94 force field. Multiple ligand docking was carried out in vina software using a customized python script. All the 75 unique ligands from pharmacophore based virtual screening were ranked according to their binding affinities against *Trypanosome brucei* Trypanothione Reductase and top two best hit selected. Maestro v14.1, ligplot and pymolv2.6 were used for visualization of the enzyme and the two ligands. To carry out *in silico* pharmacokinetic evaluation, SWISS ADME online server was used to predict the Absorption, distribution, metabolism and excretion of the top 2 ligands. Moreover, pharmacokinetic screening was also done using pkCSM, StopTox and Pro-Tox II online webservers.

### 3. All atoms Molecular Dynamics simulations and Free binding energy calculation

The protein-ligand system stabilities were performed using charmm36-jul2022 force field. Compatible topology files were created using Networkx v2.3, Numpy and a customized python script. Molecular dynamics simulation of all atoms was carried out using Groningen Molecular Simulation package (GROMACS), version 2024.4. A dodecahedron box was defined with a cut off of 1.0 Å from all the molecular edges. Simple point charge (spc216) water model was used to solvate the system and neutralization achieved by addition of Either sodium or chloride ions. For optimal structural geometry, energy minimization was carried out using steepest descent minimization integrator at 0.01nm energy step size.

The maximum number of energy minimization steps was set at 50,000 and the system fixed to attain a maximum force of less than 10.0 kJ/mol. The system was equilibrated in a two-step canonical ensemble approach, with each equilibration step set at 100 picoseconds. The first step involved Number of particles, volume and Temperature (NVT), and it was performed at 300K using Berendsen Temperature coupling. The second equilibration step included the Number of particles, pressure and Temperature (NPT), and it was carried out at 1 atmosphere in all molecule directions at a constant temperature of 300K using the Parrinello-Rahman as the Barostat algorithm. The particle Mesh Ewald (PME) coulomb type was utilized for the ensembles processes involving long range electrostatics interactions and the gap cut off set at 8.0 Å.

To constraint the bonds between all the system atoms, the LINCS algorithm was utilized. The equilibrated system was then set for 100 nanoseconds of all atoms Molecular Dynamics with integration of 2fs. The molecular dynamics simulations were carried out using ZUPUTO High performance computing, which is hosted by the West Africa Center for Cell Biology of Infectious Pathogens, using 7212 X NVIDIA TESLA P100 16GB and the coordinates updated every 100 Picoseconds. This Trajectory production runs produced 1000 simulation frames, which were thereafter stripped off the periodic boundary conditions and aligned with starting molecules as the reference structures. Comparative analyses for the global and conformational changes were done using GROMACS in built packages. The analysis included Root mean square Deviation (RMSD), Root Mean Square Fluctuation (RMSF), Radius of Gyration (RG), solvent accessible surface area (SASA) and number of Hydrogen Bonds (H-Bond) for the 2 protein-ligand complexes.

To estimate the free binding energy of the protein-ligand complexes, Molecular Mechanics Poisson-Boltzmann Surface Area (MMPBSA) method was used. To calculate the binding free energy Δ**G_bind_** of the receptor-ligand complex, the following equation was employed.

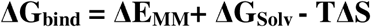

Where,

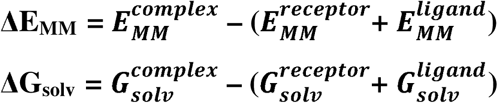

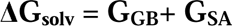

**T**Δ**S** is the free energy entropic contribution and was ignored during computation.

The energy terms (complex, receptor and ligand) were first extracted using gmx_MMPBSA version 1.6.3. The energy differences between the complex and the sum of energies of the receptor and ligand for each term were then computed. Finally, the summation of the terms was done, which included the combination of the molecular mechanic’s energy differences and the solvation energy differences. The total binding free energy was obtained using last 100 molecular dynamics simulation frames of the receptor-ligand complex.

## Results

### Pharmacophore querry modelling and validation

**Fig 1.1.**
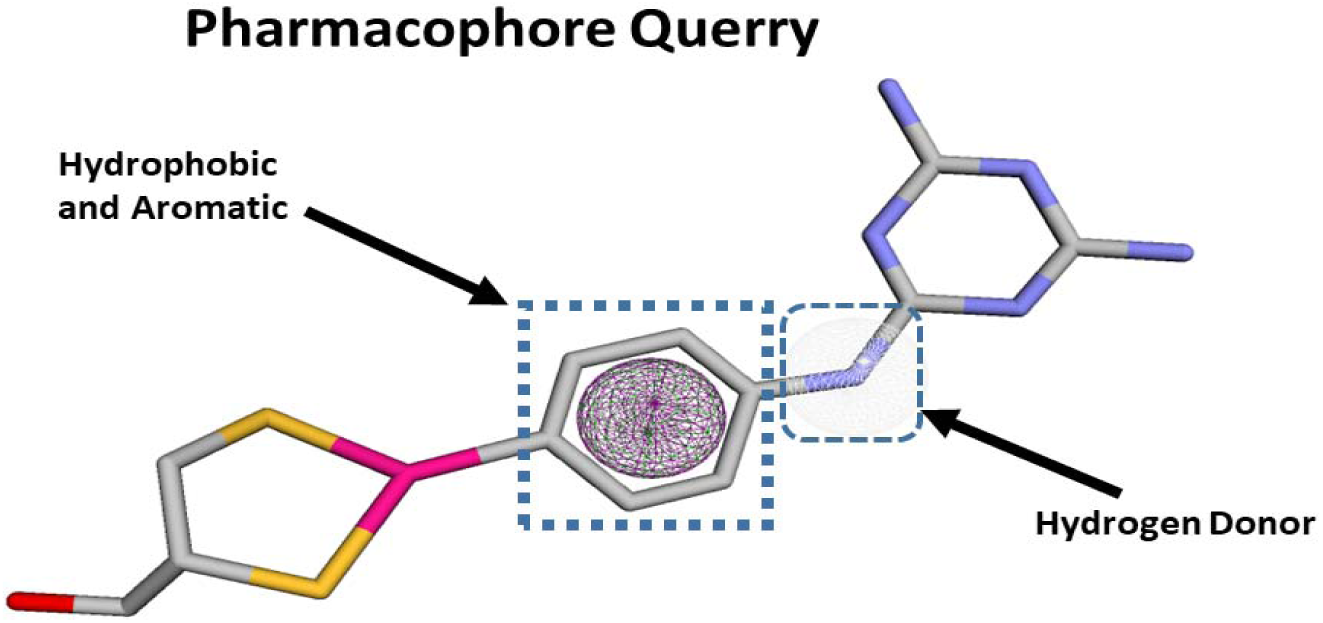
Pharmacophore querry derived from Melarsoprol. The model incorporates 3 pharmacophore properties i.e. Hydrophobic, aromatic and hydrogen donor.

**Fig 1.2.**
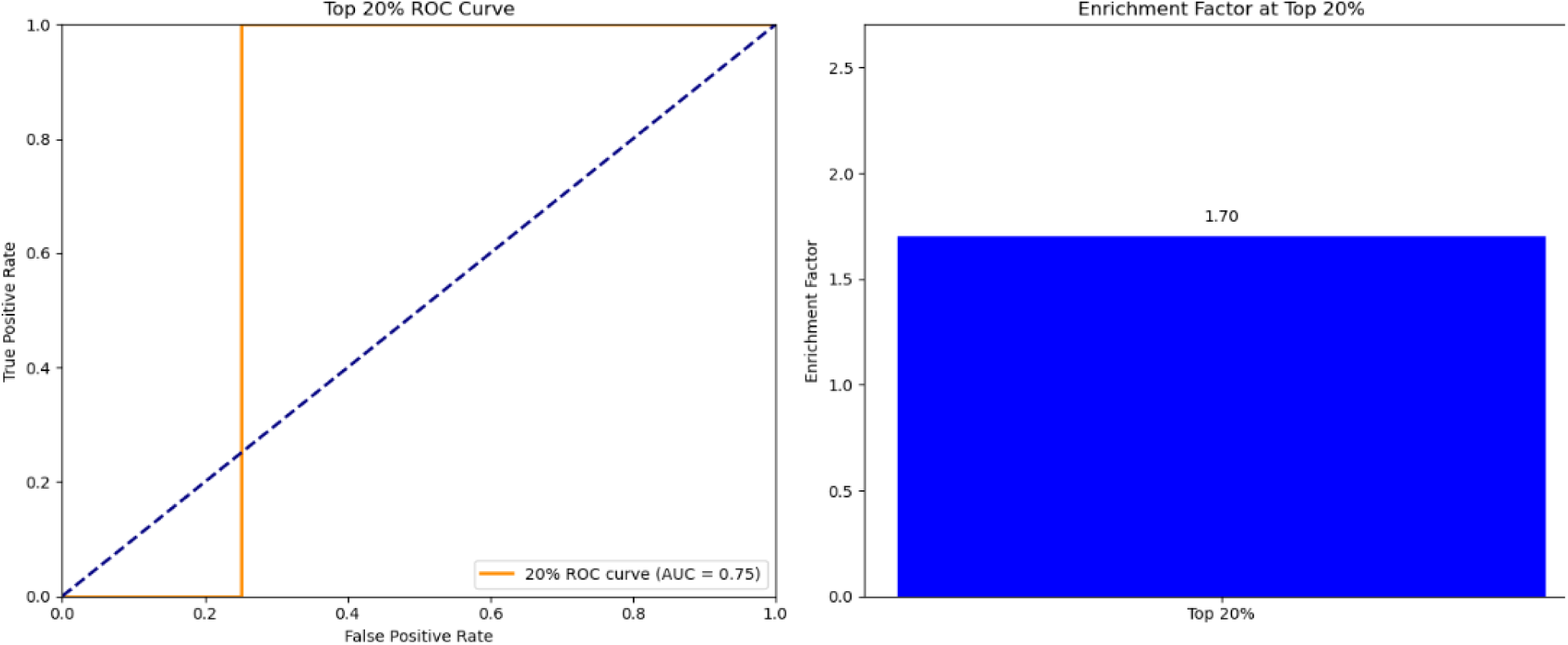
pharmacophore querry validation using Receiver operating characteristics curve. The top 20% of actives and decoys library has AUC of 0.75 and 1.70 enrichment factor.

### Virtual screening, multiple ligand docking and protein-ligand interaction analysis

**Table 1:**
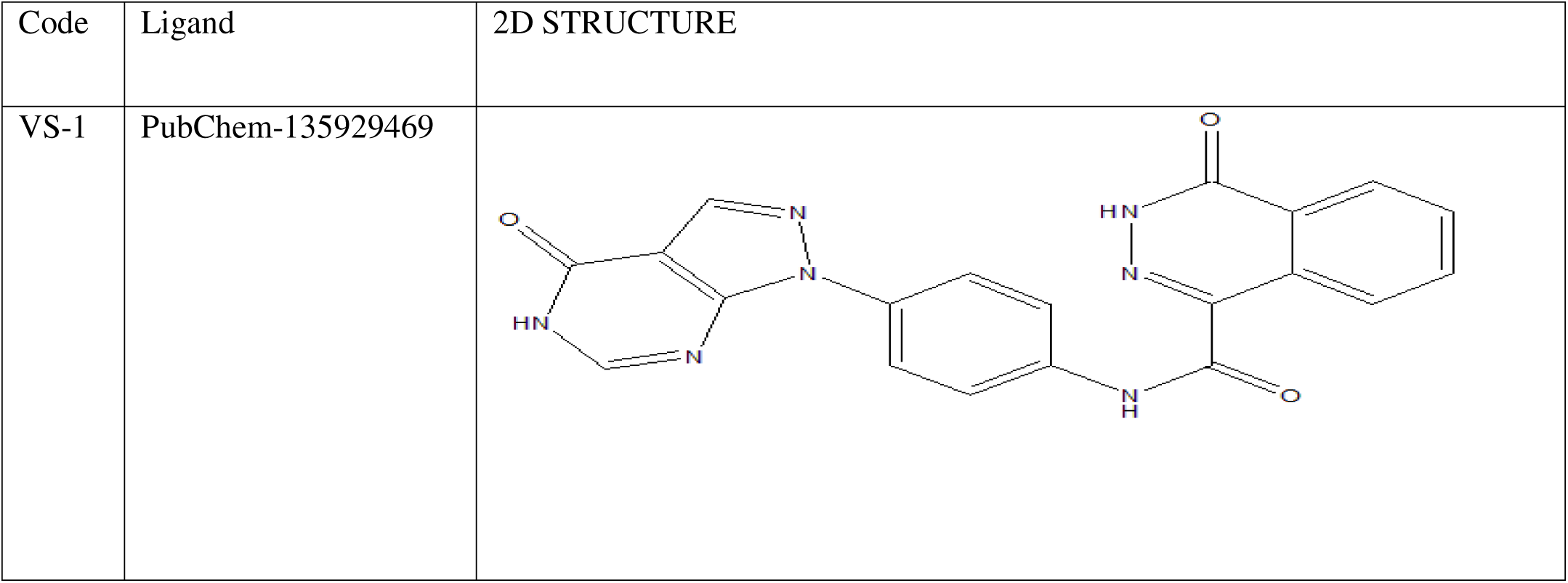
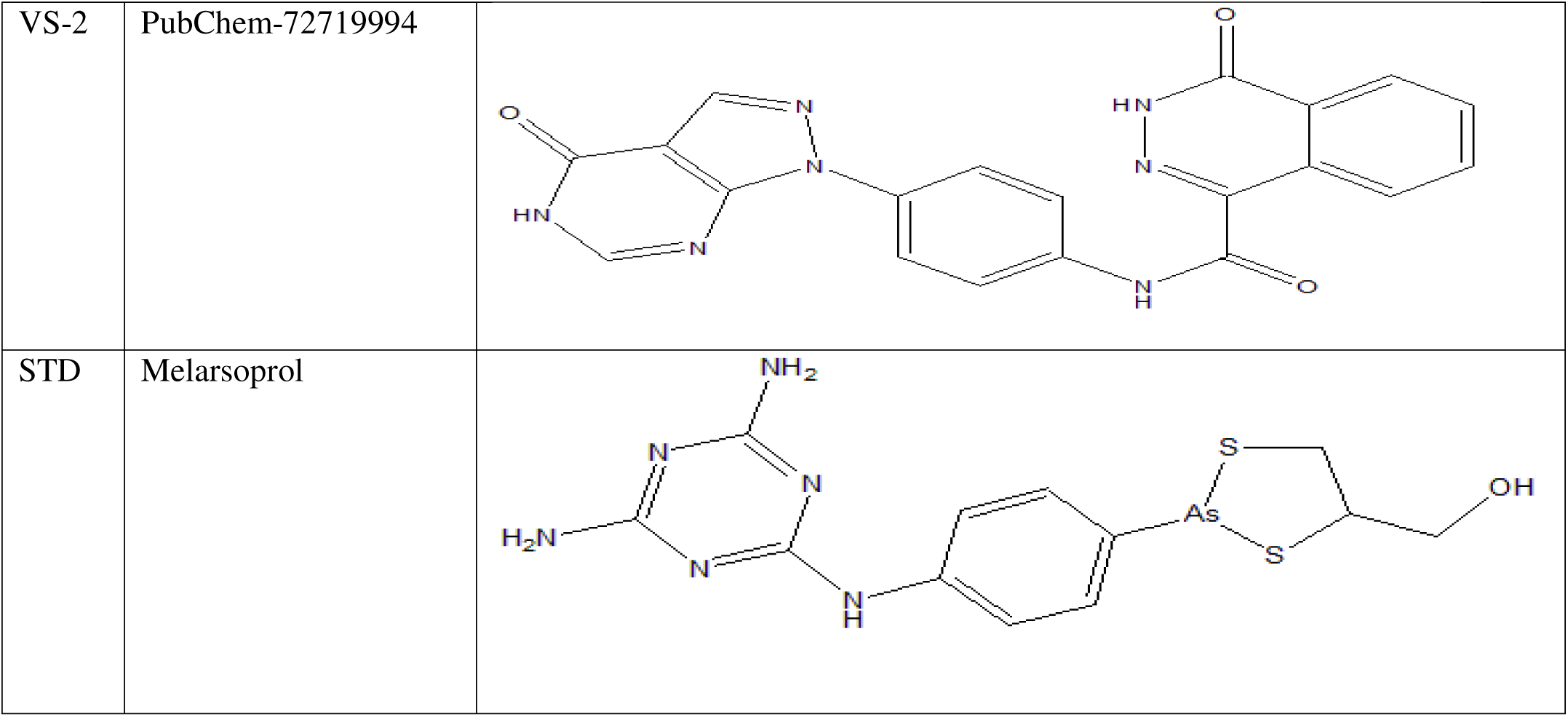
2D structures of the top two ligands obtained from structure-based virtual screening and standard drug (melarsoprol).

| Code | Ligand | 2D STRUCTURE |
| --- | --- | --- |
| VS-1 | PubChem-135929469 |  |
| VS-2 | PubChem-72719994 |  |
| STD | Melarsoprol |  |

**Table 1:** Docking Scores of the top two ligands against *T.b brucei* Trypanothione Reductase.

| Code | Docking score | Name of the compound |
| --- | --- | --- |
| VS-1 | -11.3<br>Kcal/Mol | 4-oxo-N-[4-(4-oxo-5H-pyrazolo[3,4-d]pyrimidin-1-yl)phenyl]-3H-phthalazine-1-carboxamide |
| VS-2 | -10.6<br>Kcal/Mol | 3-(benzo[d][1,3]dioxol-5-yl)-N-(4-oxo-5,6,7,8-tetrahydro-4H-thiazolo[5,4-c]azepin-2-yl)-1H-pyrazole-5-carboxamide |
| STD | -7.8<br>Kcal/Mol | Melarsoprol |

**Figure 2.1:**
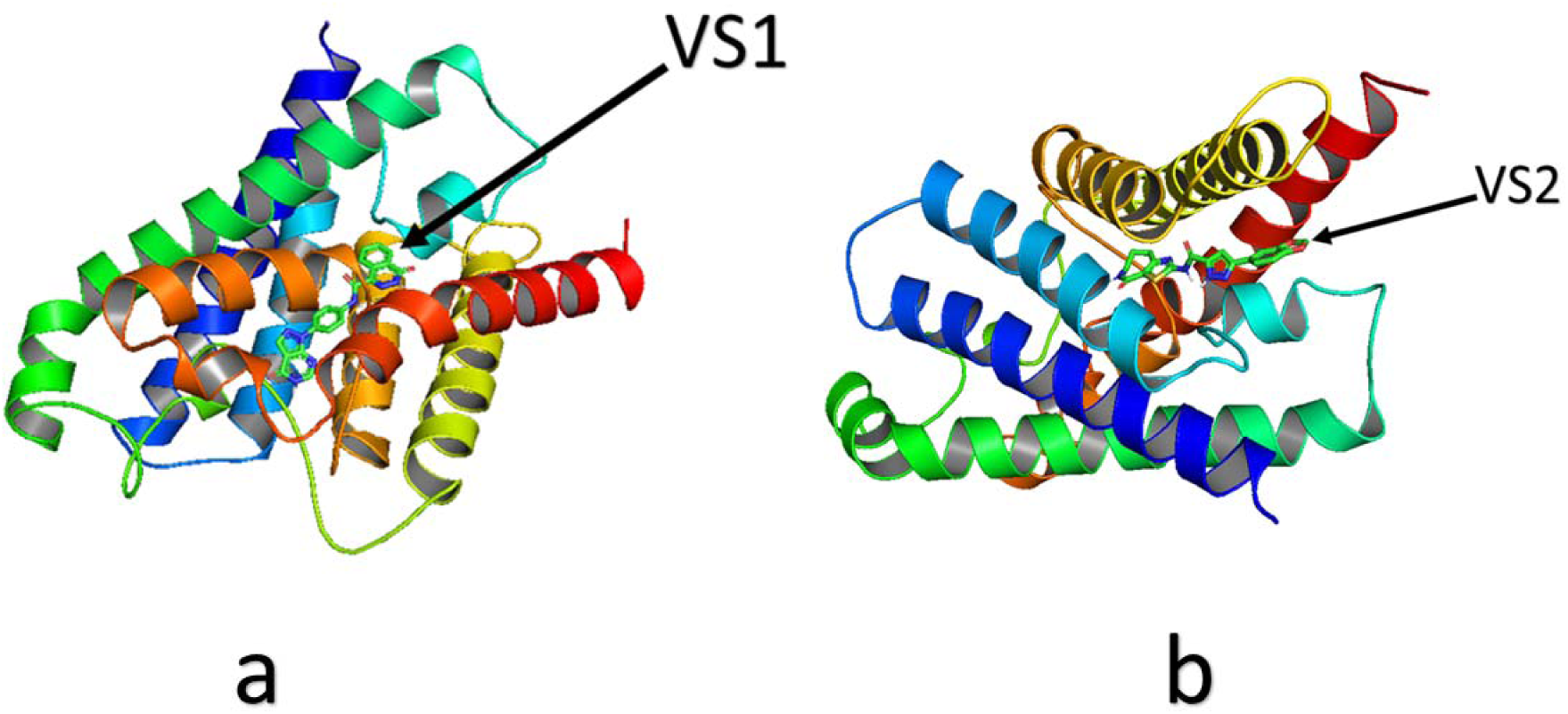
*Trypanosome brucei* Trypanothione reductase enzyme superimposed image with top 2 ligands i.e. VS1 (a) and VS2 (b).

**Fig 2.2.**
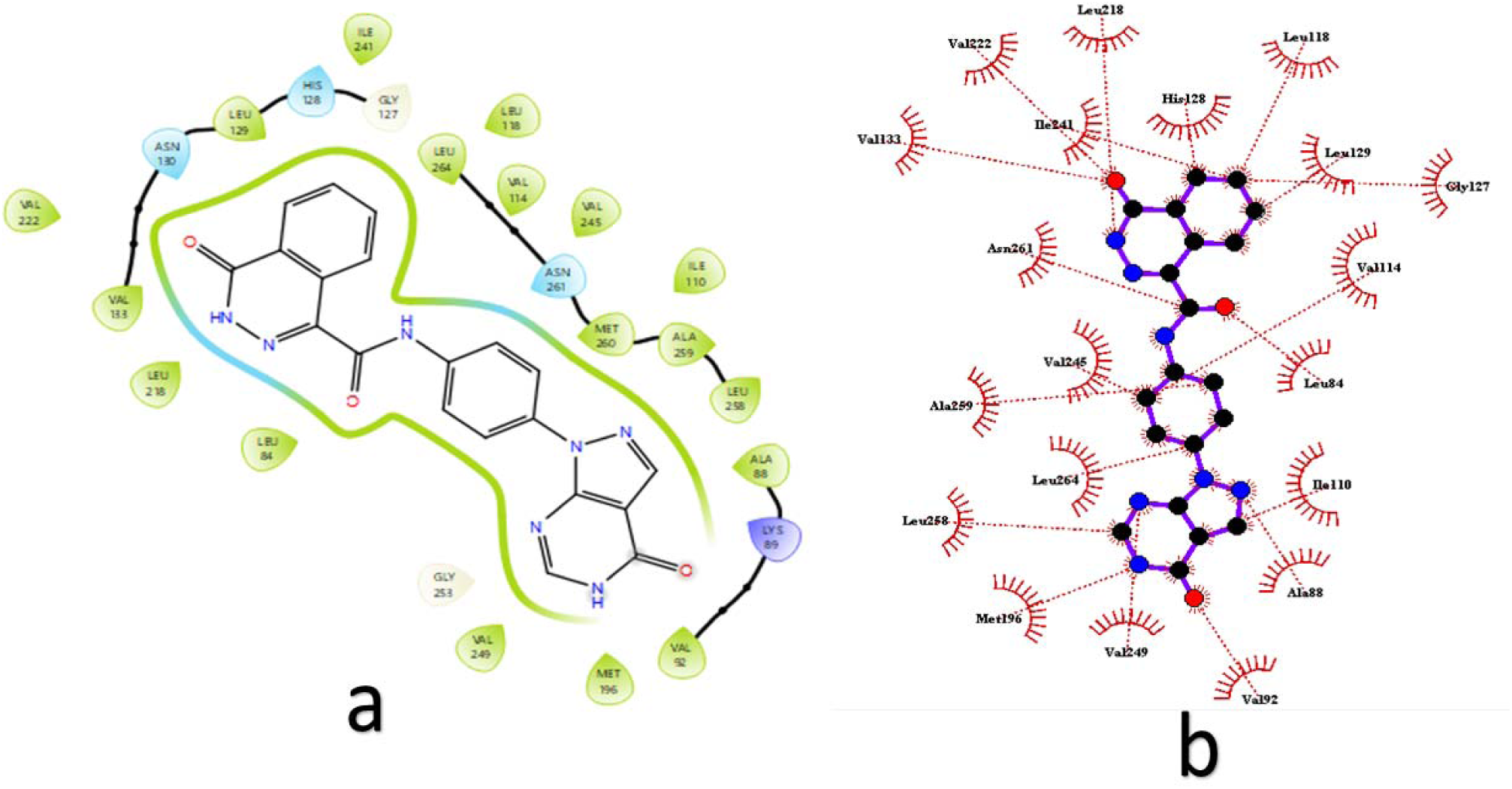
(a). 2D pose view interaction diagram between VS1 and *T.b* TR. The ligand is held in the enzyme’s active site by hydrophobic interactions facilitated by nearby amino acids such as LEU 218, ASN 130, MET 260, and ILE 110. Fig 2.2 (b) shows an estimated distance of the nearby amino acids during the VS1-*T.b* TR interaction.

**Fig 2.3.**
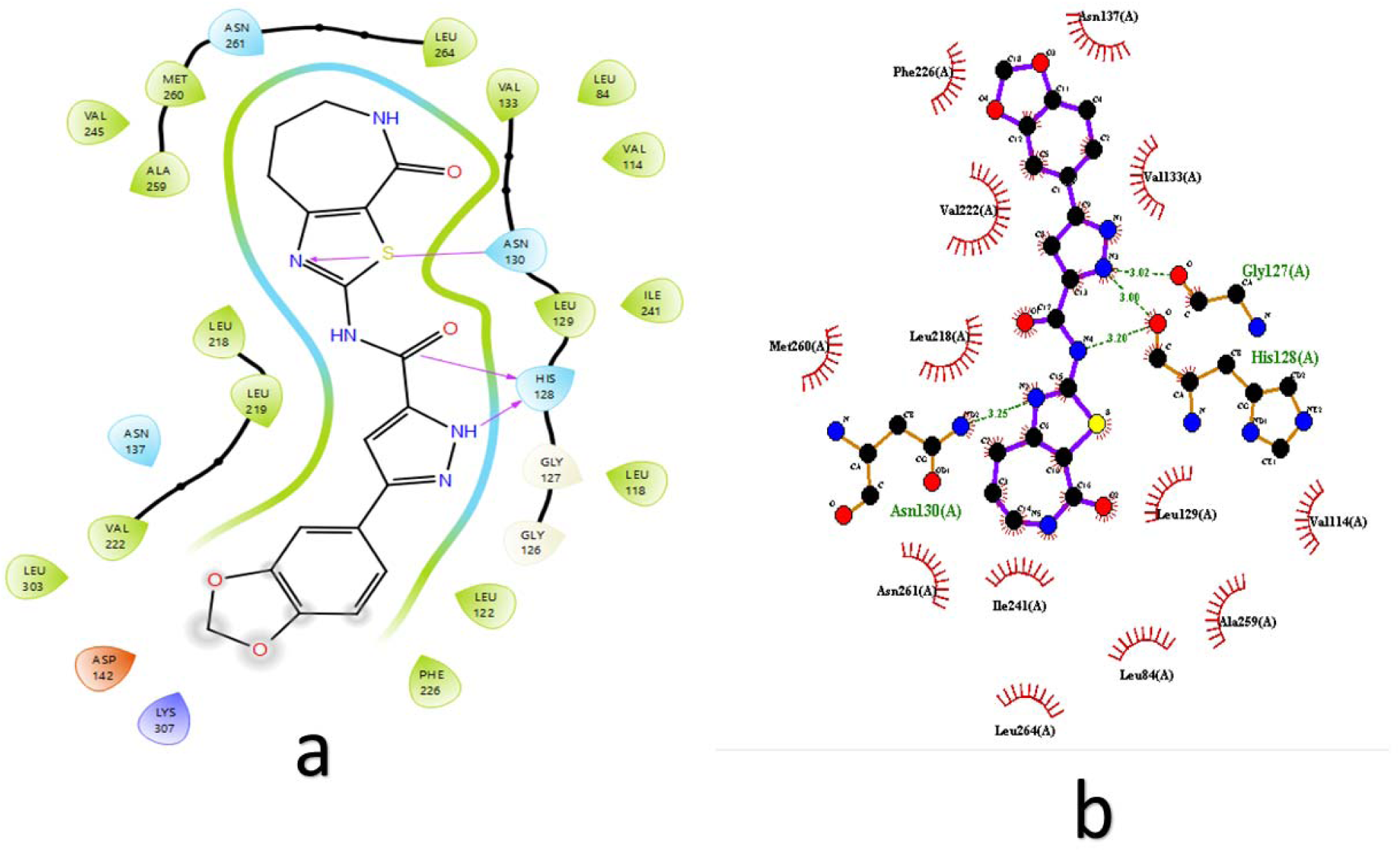
(a). 2D pose view interaction diagram between VS2 and *T.b* TR showing HIS 128 and ASN 130 as the most important amino acids in the interaction since they contribute to the formation of hydrogen bonds between the ligand and the enzyme. Fig 2.3 (b) depicts the spatial distribution of atoms in the VS2-*T.b* TR active site. The amino acid ASN 130, HIS 128 and GLY 127 are within the distance of 3.25 Å, 3.20/3.00Ä and 3.02Å respectively.

### Pharmacokinetic evaluation

**Fig 2.4:**
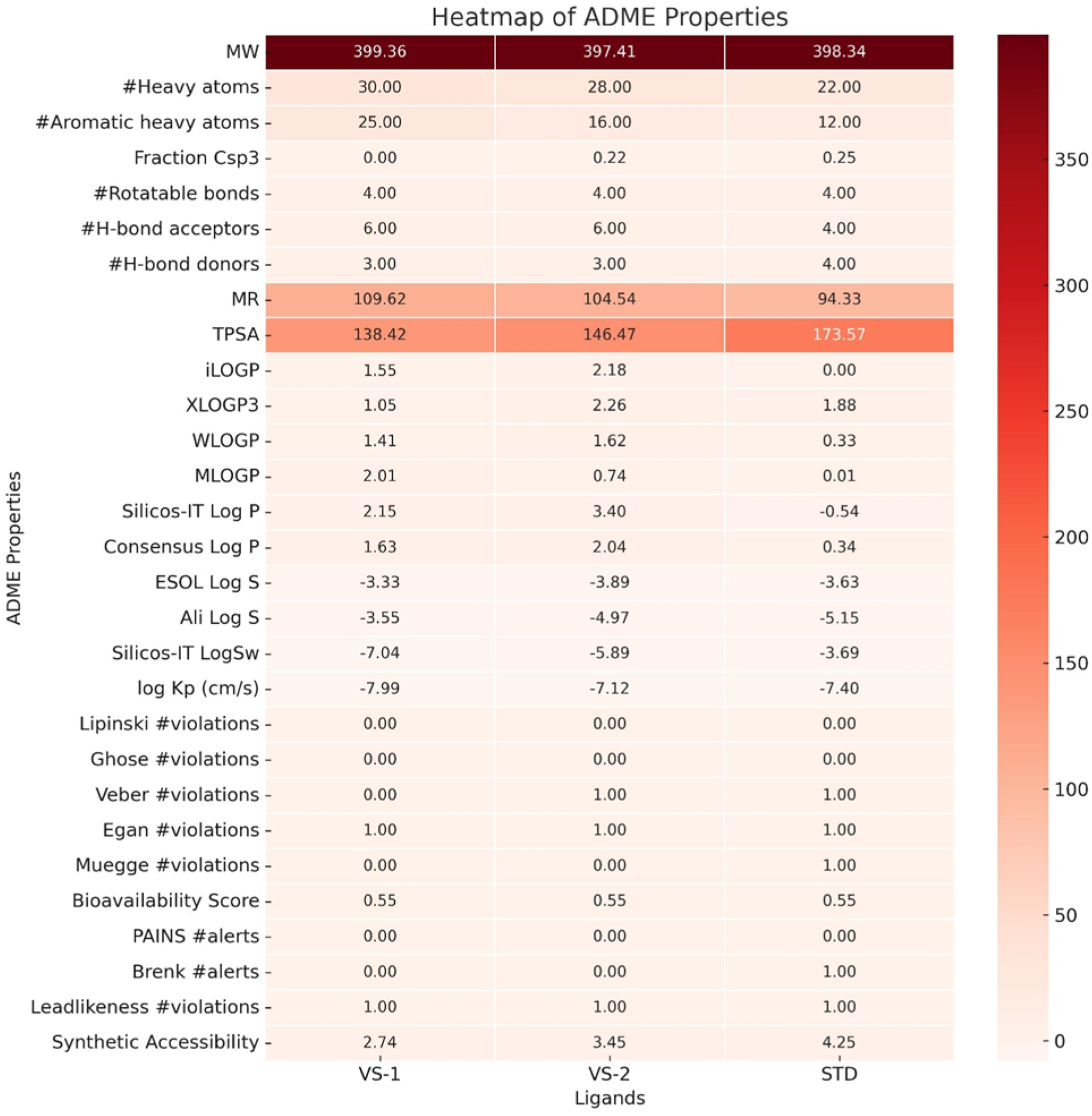
Heatmap of ADME analysis of VS1 and VS2 and the standard (drug melarsoprol) as predicted by SWISS ADME online server.

**Table 2:** pkCSM pharmacokinetic parameters of VS-1, VS-2 and STD.

| Pharmacokinetic properties |  | Ligands |  |  |
| --- | --- | --- | --- | --- |
| Properties | Model Name | VS-1 | VS-2 | STD |
| Absorption | Water solubility | -2.983 | -3.212 | -3.032 |
|  | Caco2 permeability | 0.607 | 1.183 | -0.011 |
|  | Intestinal absorption (human) | 66.799 | 86.584 | 62.349 |
|  | Skin Permeability | -2.735 | -2.736 | -2.735 |
|  | P-glycoprotein substrate | Yes | Yes | No |
|  | P-glycoprotein I inhibitor | No | No | No |
|  | P-glycoprotein II inhibitor | No | No | No |
| Distribution | VDss (human) | -0.284 | 0.46 | 1.025 |
|  | Fraction unbound (human) | 0.025 | 0.118 | 0.477 |
|  | BBB permeability | -1.677 | -1.287 | -1.016 |
|  | CNS permeability | -3.72 | -3.578 | -3.248 |
| Metabolism | CYP2D6 substrate | No | No | No |
|  | CYP3A4 substrate | Yes | No | No |
|  | CYP1A2 inhibitor | No | Yes | No |
|  | CYP2C19 inhibitor | No | No | No |
|  | CYP2C9 inhibitor | No | No | No |
|  | CYP2D6 inhibitor | No | No | No |
|  | CYP3A4 inhibitor | No | No | No |
| Excretion | Total Clearance | 0.198 | -0.132 | -0.378 |
|  | Renal OCT2 substrate | No | No | No |
| Toxicity | AMES toxicity | No | No | No |
|  | Max. tolerated dose (human) | 0.69 | 0.584 | 0.559 |
|  | hERG I inhibitor | No | No | No |
|  | hERG II inhibitor | Yes | Yes | No |
|  | Oral Rat Acute Toxicity (LD50) | 3.073 | 2.392 | 2.847 |
|  | Oral Rat Chronic Toxicity (LOAEL) | 2.358 | 1.448 | 2.577 |
|  | Skin Sensitization | No | No | No |
|  | T.Pyriformis toxicity | 0.285 | 0.289 | 0.304 |
|  | Minnow toxicity | 1.588 | 3.015 | 2.235 |

**Table 3:** Pro-Tox II toxicological parameters of VS-1, VS-2 and STD ligands.

| Classification | Target | VS-1 |  | VS-2 |  | STD |  |
| --- | --- | --- | --- | --- | --- | --- | --- |
|  |  | Prediction | Probability | Predicted | Probability | Predicted | Probability |
| Organ Toxicity | Hepatotoxicity | Active | 0.60 | Active | 0.53 | Active | 0.69 |
|  | Neurotoxicity | Active | 0.75 | Active | 0.57 | Active | 0.87 |
|  | Nephrotoxicity | Active | 0.52 | Active | 0.53 | Inactive | 0.90 |
|  | Respiratory Toxicity | Active | 0.60 | Active | 0.78 | Active | 0.98 |
|  | Cardiotoxicity | Inactive | 0.91 | Inactive | 0.61 | Inactive | 0.77 |
| Toxicity end points | Carcinogenicity | Active | 0.56 | Inactive | 0.51 | Inactive | 0.62 |
|  | Immunotoxicity | Inactive | 0.99 | Inactive | 0.98 | Active | 0.96 |
|  | Mutagenicity | Inactive | 0.61 | Active | 0.51 | Inactive | 0.97 |
|  | Cytotoxicity | Inactive | 0.86 | Inactive | 0.66 | Inactive | 0.93 |
|  | BBB-barrier | Active | 0.86 | Inactive | 0.50 | Inactive | 1.0 |
|  | Clinical Toxicity | Active | 0.61 | Active | 0.56 | Inactive | 0.56 |
| Tox21-Nuclear receptor signaling pathways | AhR | Inactive | 0.61 | Inactive | 0.78 | Inactive | 0.97 |
|  | AR | Inactive | 0.98 | Inactive | 0.93 | Inactive | 0.99 |
|  | AR-LBD | Inactive | 0.99 | Inactive | 0.97 | Inactive | 0.99 |
|  | Aromatase | Inactive | 0.87 | Inactive | 0.94 | Active | 1.0 |
|  | ER | Inactive | 0.91 | Inactive | 0.79 | Active | 0.99 |
|  | ER-LBD | Inactive | 0.99 | Inactive | 0.94 | Active | 1.0 |
|  | PPAR-Gamma | Inactive | 0.97 | Inactive | 0.94 | Inactive | 0.99 |
|  | nrf2/ARE | Inactive | 0.95 | Inactive | 0.94 | Inactive | 0.88 |
| Tox21-stress response pathways | HSE | Inactive | 0.95 | Inactive | 0.94 | Inactive | 0.88 |
|  | MMP | Inactive | 0.76 | Inactive | 0.81 | Inactive | 0.70 |
|  | P53 | Inactive | 0.92 | Inactive | 0.85 | Inactive | 0.96 |
|  | ATAD5 | Inactive | 0.82 | Inactive | 0.89 | Inactive | 0.99 |

AhR: Aryl hydrocarbon Receptor, AR: Androgen Receptor, ER: Estrogen Receptor Alpha, ER-LBD: Estrogen Receptor Ligand Binding Domain, PPAR-Gamma: Peroxisome Proliferator Activated Receptor Gamma, nrf2/ARE: Nuclear factor (erythroid-derived 2)-like 2/antioxidant responsive element, HSE: Heat shock factor response element, MMP: Mitochondrial Membrane Potential, p53: Phosphoprotein (Tumor Suppressor), ATAD5: ATPase family AAA domain-containing protein 5.

### Molecular dynamics simulations

**Fig 3.1.**
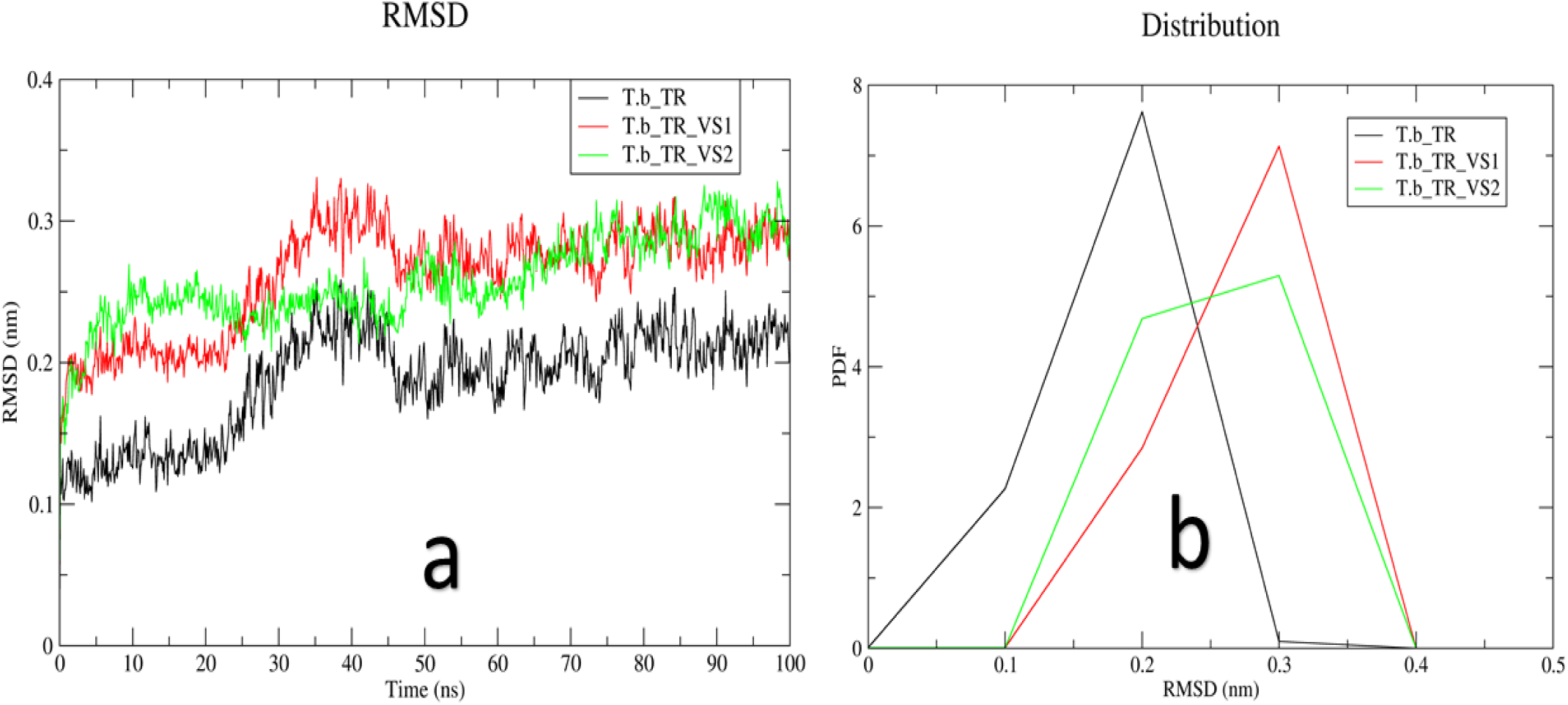
(a): Root mean square deviation of *Trypanosome brucei* Trypanothione Reductase enzyme (Apo form) and Holo form of the enzyme when complexed with VS1 and VS2 ligands in the 100 ns simulation time. Fig 3.1 (b) shows the probability density function of the RMSDs across 100 ns.

**Fig 3.2.**
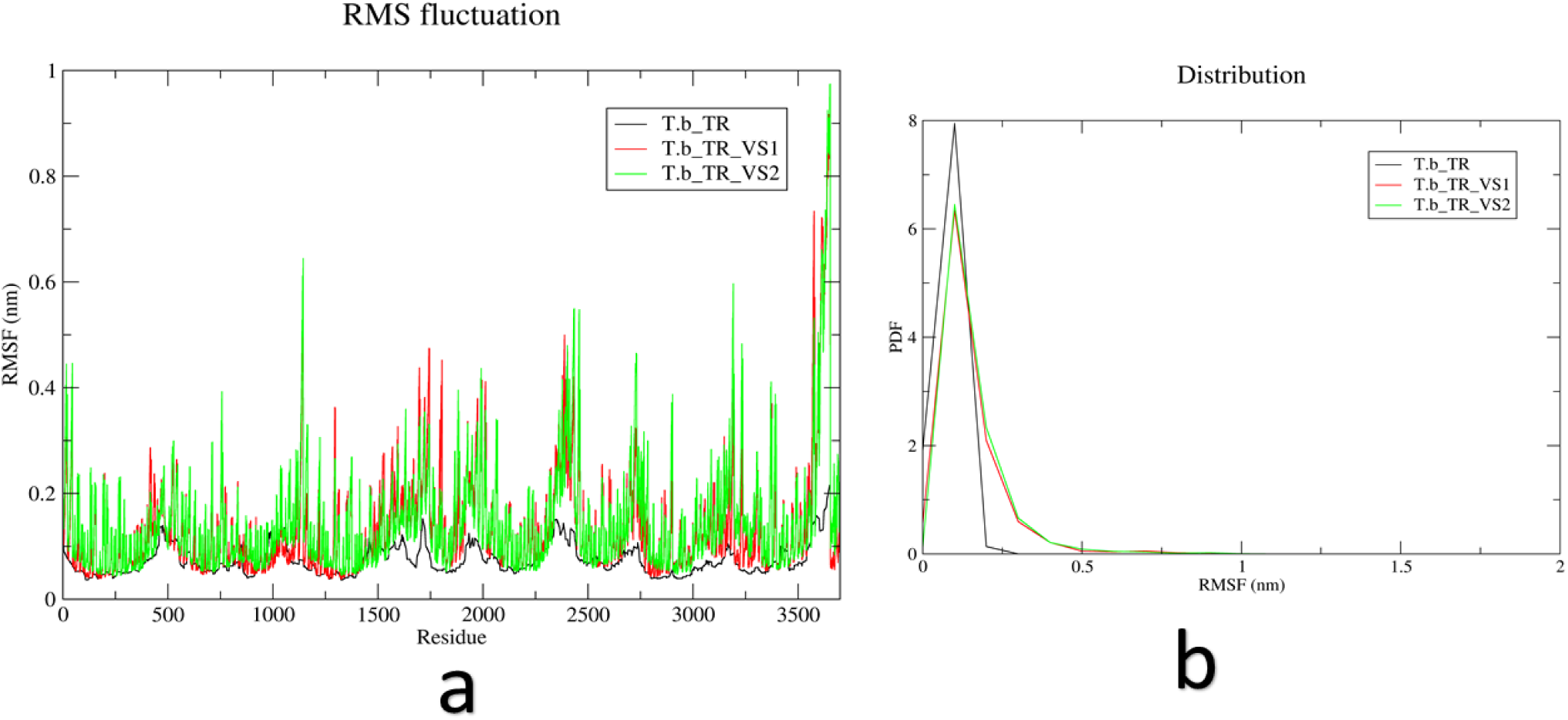
(a): Root mean square fluctuation plot for unbound *T.b* TR and *T.b* TR complexed with VS1 and VS2 during the 100 ns all atoms molecular dynamics simulation. Fig 3.2 (b) Shows the probability density function of the RMSFs during the 100 ns simulation time.

**Fig 3.3.**
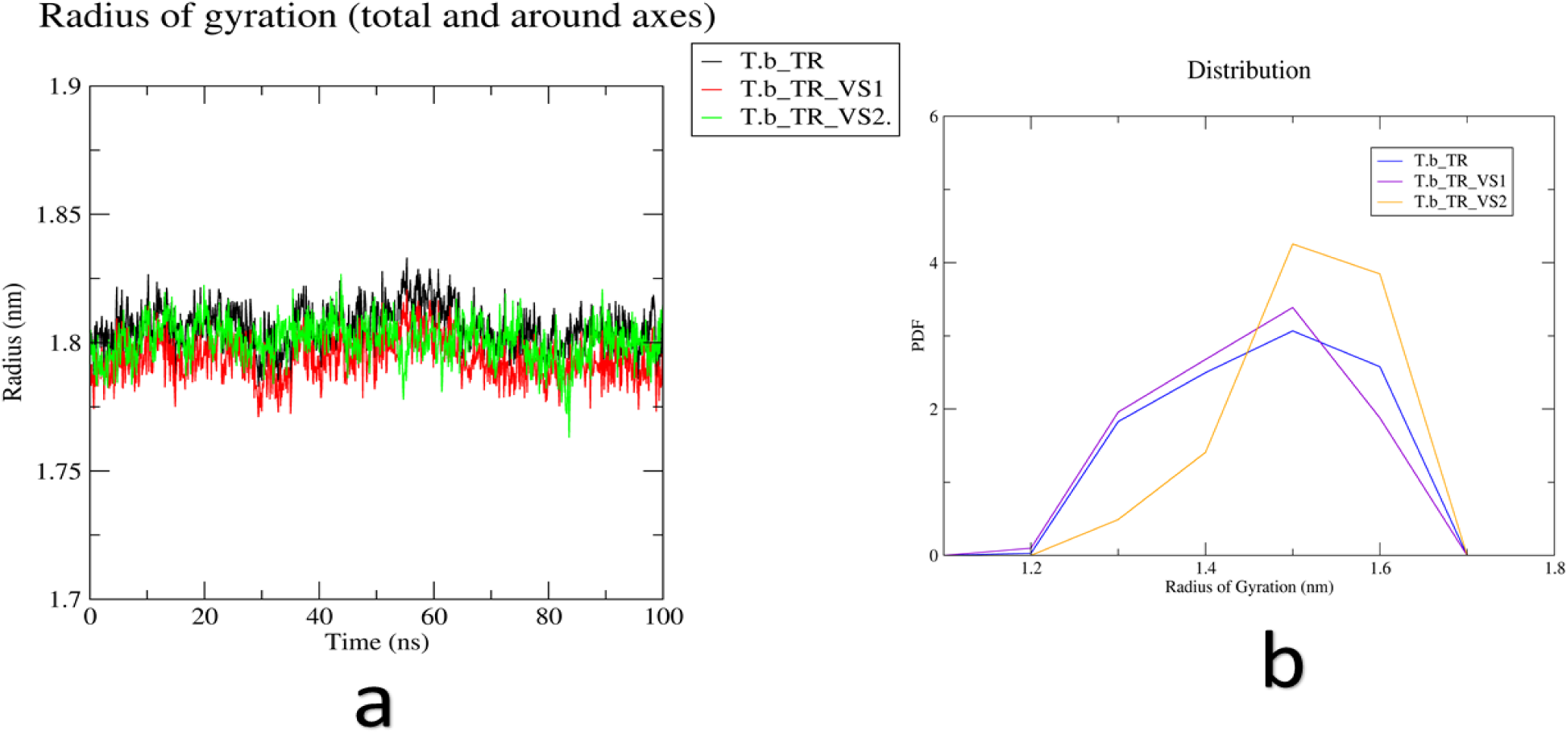
(a). Radius of gyration analysis of Unbound *T.b* TR and VS1 and VS2 bound *T.b* TR during the 100 ns of molecular dynamics simulation. Fig 3.3 (b) shows probability density function of RGs in 100 ns the simulation trajectories.

**Fig 3.4:**
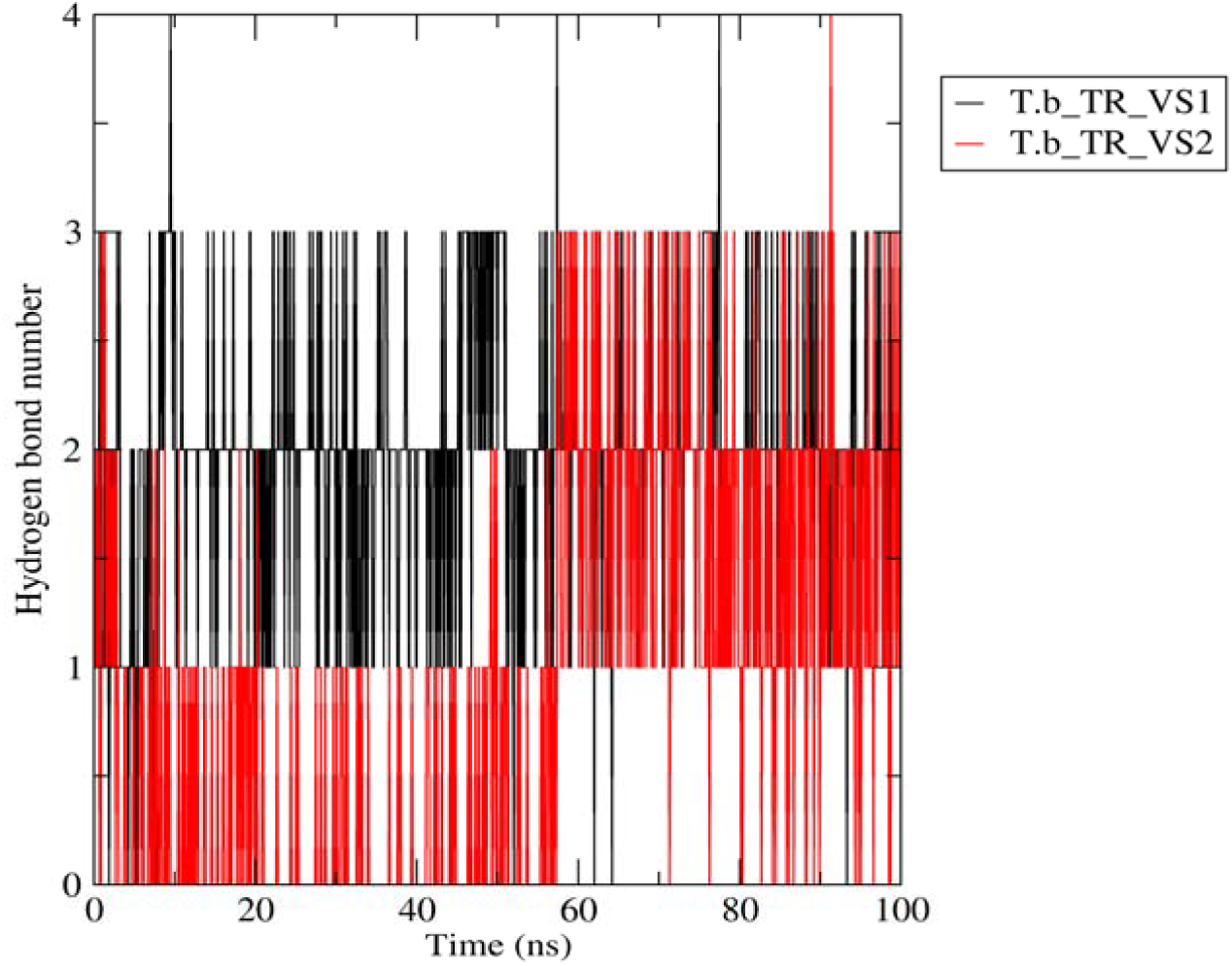
Number of hydrogen bond interactions between *T.b* TR and ligands VS1 and VS2 during 100 ns simulation time.

**Fig 3.5:**
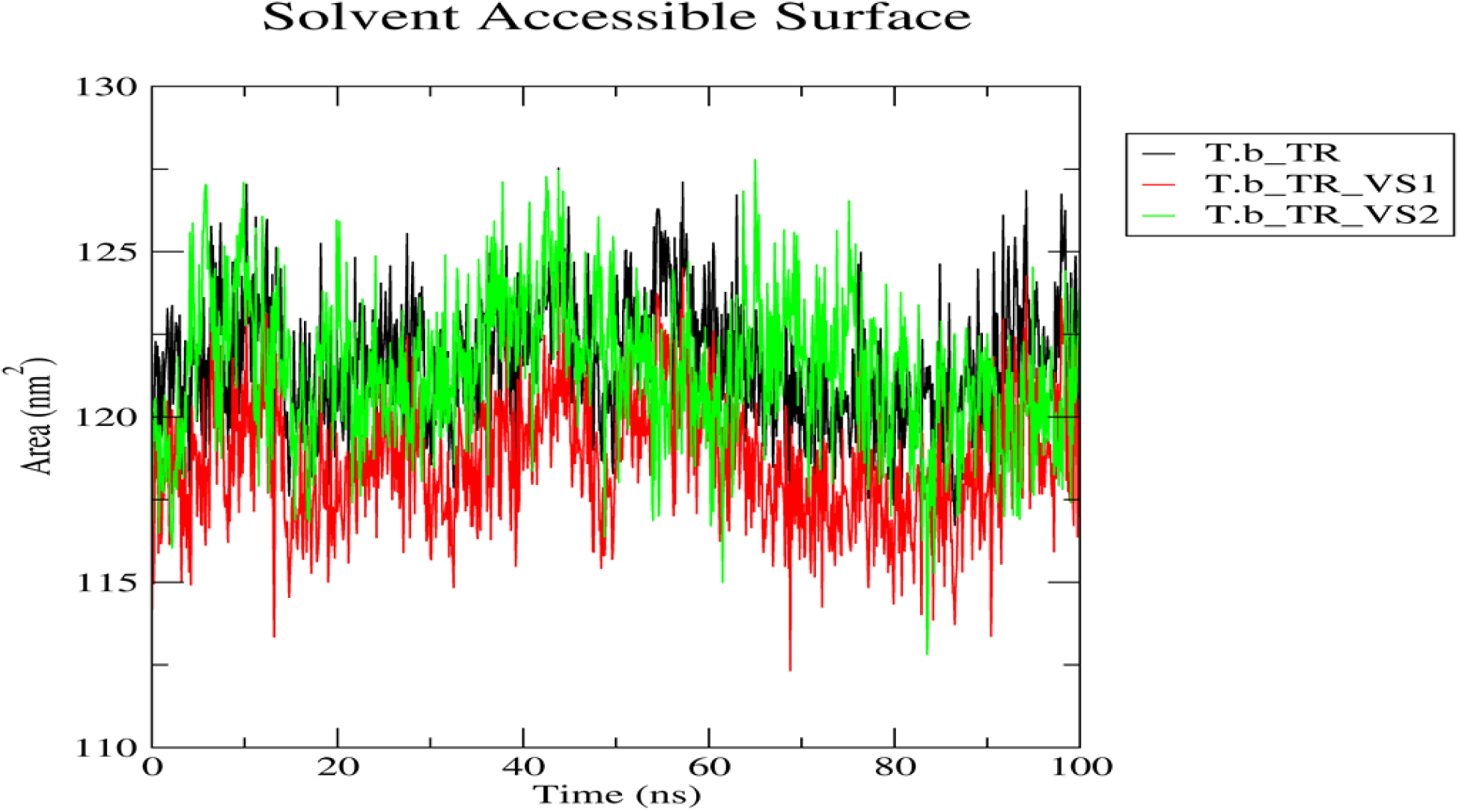
solvent accessible surface area analysis of the unbound *T.b* TR and the bound form of *T.b* TR with ligands VS1 and VS2 across 100 simulation time.

**Fig 3.6.**
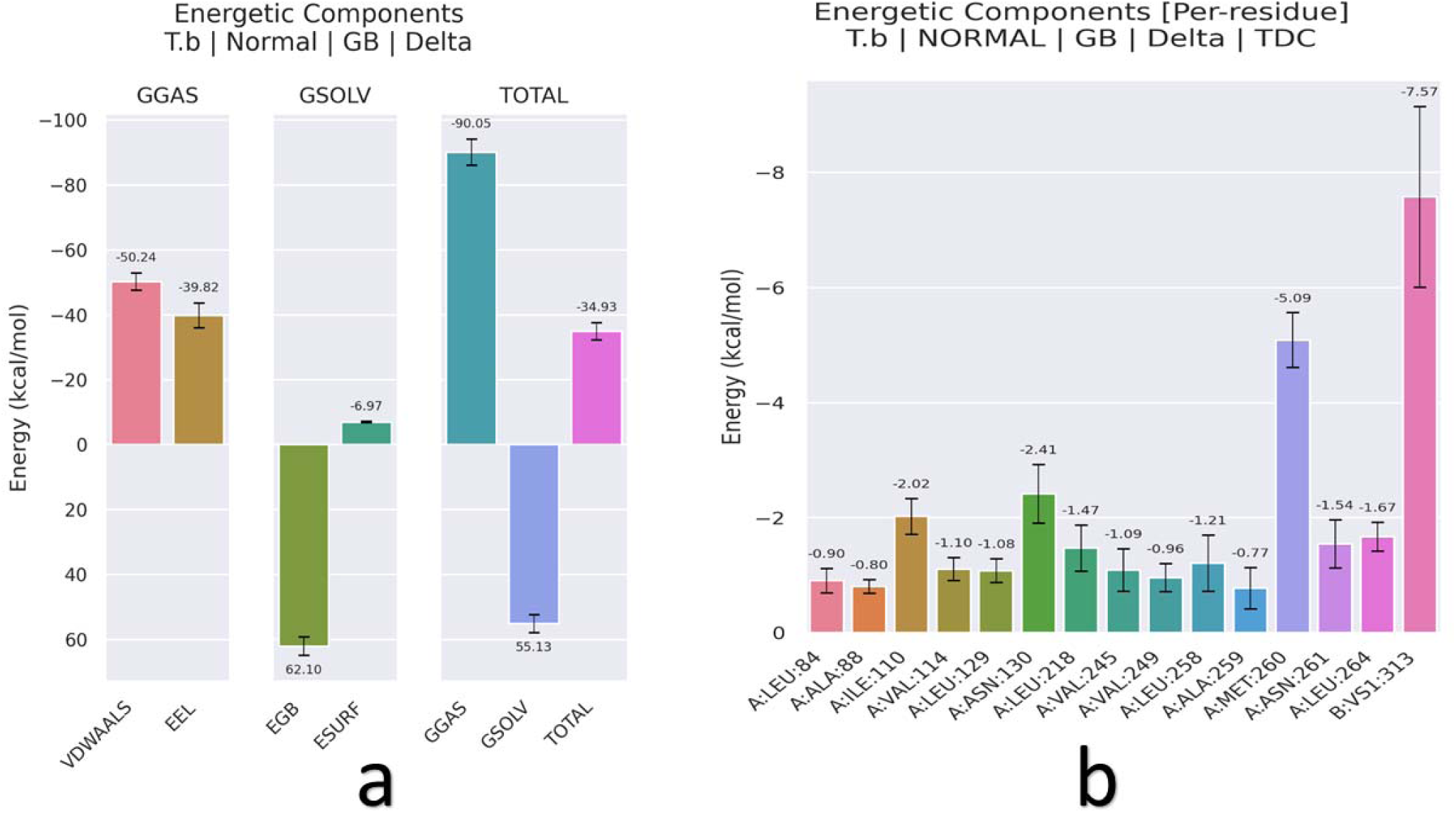
(a). Binding free energy estimation for *T.b* TR-VS1 complex as computed using gmx-MMPBSA tool v1.6.3. The free binding energy for the receptor-ligand complex was estimated using snapshots of the last 100 simulation frames and was found to be −30.93 kcal/Mol. Fig 3.6 (b) shows per residue decomposition of *T.b* TR-VS1. The ligand VS1 had contributed to the largest free binding energy (−7.57 kcal/Mol) followed by MET 260 (−5.09 kcal/mol).

**Fig 3.7.**
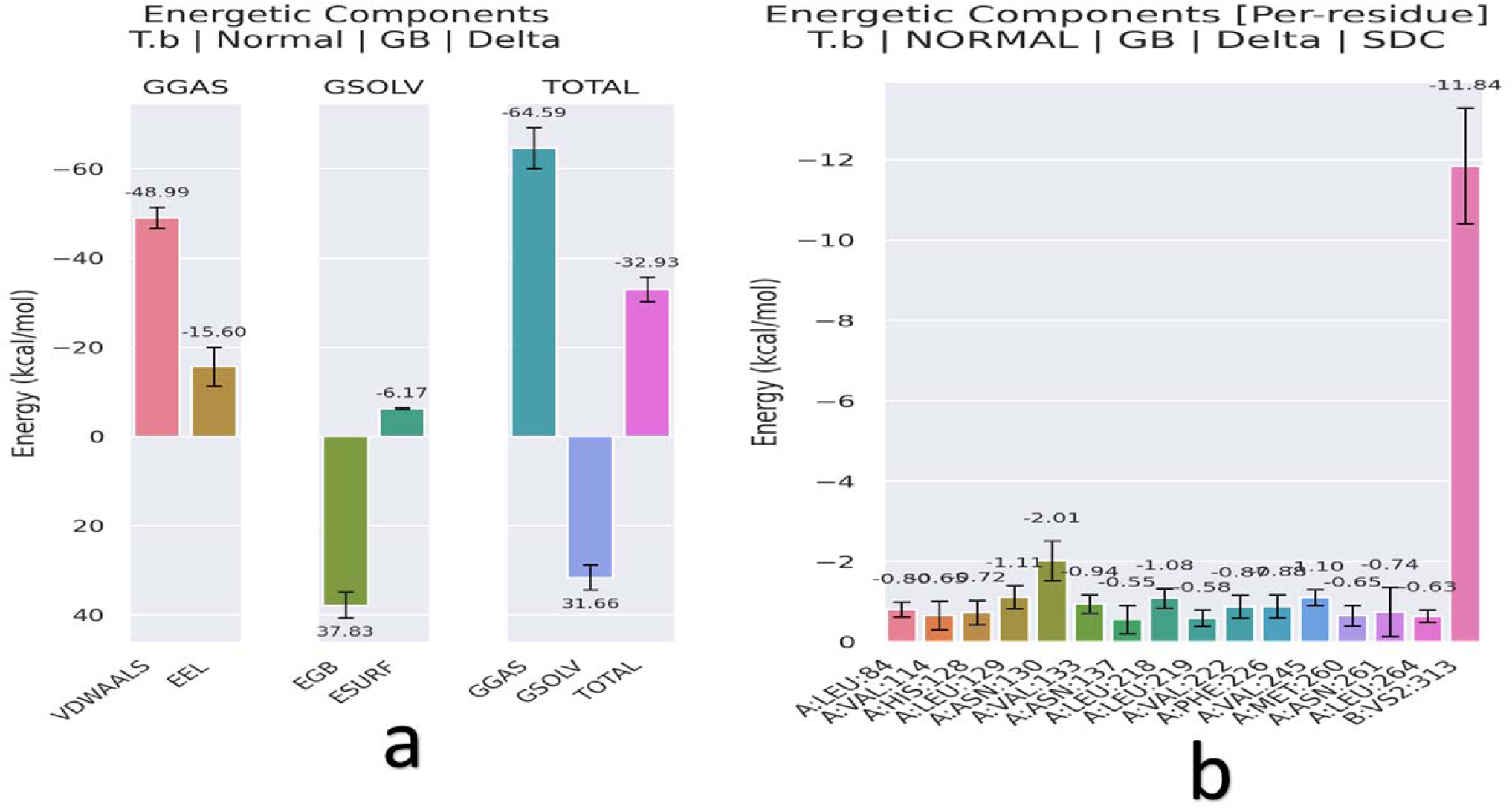
(a). Binding free energy estimation for *T.b* TR-VS2 complex as computed using gmx-MMPBSA tool v1.6.3. The free binding energy for *T.b* TR-VS2 was estimated using snapshots of the last 100 simulation frames and was found to be −32.93 kcal/Mol. Fig 3.7 (b) shows per residue decomposition of *T.b* TR-VS2. The ligand VS2 contributed to the largest free binding energy (−11.84 kcal/Mol).

## Discussion

### Pharmacophore Query Generation

In this study, Melarsoprol, a standard drug for the treatment of late-stage Human African Trypanosomiasis (HAT), was selected for pharmacophore query generation due to its known inhibitory effect on Trypanothione Reductase (*T.b* TR) [10]. Melarsoprol’ s effectiveness stems from its ability to interfere with the redox metabolism of *Trypanosome brucei*, making it a strong candidate for generating a pharmacophore model. Despite its clinical relevance, the severe side effects associated with melarsoprol, including reactive encephalopathy, and the increasing emergence of drug-resistant *Trypanosome brucei* strains necessitate the discovery of alternative therapeutic agents with enhanced safety profiles and efficacy [11]. Key pharmacophore features were derived from Melarsoprol’ s structure, including hydrophobic, aromatic, and hydrogen donor characteristics. These features were critical to capturing the binding interaction between Melarsoprol and *Trypanosome brucei* Trypanothione Reductase’s active site. The selected pharmacophore properties effectively modeled the interaction required to inhibit the enzyme [12]. However, modeling hydrophobic interactions posed some challenges due to the spatial arrangement of hydrophobic residues in the active site, requiring multiple rounds of optimization to refine the pharmacophore model.

### Validation of the Pharmacophore Model

The generated pharmacophore was validated using a two-step approach. First, the pharmacophore query was screened against a library from the ZINC database to retrieve known inhibitors of *T.b* TR. The model successfully retrieved melarsoprol from the database, confirming that the pharmacophore adequately represents the key features required for binding [13]. Further validation was conducted using a Receiver Operating Characteristic (ROC) curve and Enrichment Factor (EF) based on a dataset comprising 14 active inhibitors and 255 decoys. The ROC analysis yielded an Area Under the Curve (AUC) of 0.75, while the Enrichment Factor at the top 20% of the ranked compounds was 1.7. These results demonstrate the pharmacophore’s ability to distinguish between active and inactive compounds, confirming its reliability for virtual screening. This level of performance is comparable to previous studies in pharmacophore-based virtual screening, where AUC values above 0.70 are typically considered acceptable for robust models [13]. Furthermore, the optimization of the query, such as the number of hydrogen bond donors and acceptors relative to melarsoprol and ensuring the molecular weight remained within the specified range, improved the model’s ability to retrieve active compounds from the database.

### Structure-Based Virtual Screening and pharmacokinetics evaluation

Using the validated pharmacophore model, structure-based virtual screening was performed across six compound libraries, including PubChem, ZINC, MCULE, Enamine, and Coconut database [14]. The optimized pharmacophore query was used to screen 56,184,750 unique compounds. The top 15 hits from each database were selected for further evaluation based on their binding affinity to *T.b* TR. The top two compounds, VS-1 and VS-2, were identified based on their docking scores (−11.3 kcal/Mol and −10.6 kcal/Mol, respectively), which were significantly better than Melarsoprol (−7.8 kcal/Mol). This suggests that these compounds may exhibit stronger binding interactions with the enzyme’s active site. Both compounds demonstrated favorable interactions with key residues in the active site, including ILE 110, ASN 130, and MET 260 for VS-1 and HIS 128 and ASN 130 for VS-2, as visualized in the docking analysis. These interactions likely contribute to the enhanced binding affinity compared to the standard drug. Additionally, ADME analysis of the top two compounds revealed promising pharmacokinetic properties, including high a better Blood Brain Barrier permeability and moderate toxicity end points. Both VS-1 and VS-2 were predicted to have lower organ toxicity in relation to hepatotoxicity, neurotoxicity, respiratory toxicity and cardiotoxicity risks compared to Melarsoprol, making them suitable candidates for further investigation.

### All-Atom Molecular Dynamics Simulation

The stability and dynamic behavior of the *Trypanosome brucei* Trypanothione Reductase (*T.b* TR) enzyme in complex with the top two ligands (VS-1 and VS-2) were assessed using all-atom molecular dynamics (MD) simulations for 100 nanoseconds using Gromacs v 2024.4 [15]. The goal of the simulations was to evaluate the conformational stability of the protein-ligand complexes and to understand the molecular interactions that govern binding affinity.

The root mean square deviation (RMSD) analysis indicated that both complexes reached equilibrium within the first 20 ns, with minimal fluctuations thereafter (Figure 3.1a). This suggests that both VS-1 and VS-2 formed stable complexes with *T.b* TR over the course of the simulation. Notably, the RMSD of the *T.b* TR in the unbound form showed higher fluctuations compared to the ligand-bound forms, highlighting the stabilizing effect of ligand binding on the enzyme’s structure [16]. The probability density function of the RMSD values (Figure 3.1b) further supports this observation, as both complexes maintained low and stable RMSD distributions throughout the simulation.

The root mean square fluctuation (RMSF) plot (Figure 3.2a) provided insights into the flexibility of specific amino acid residues. Both ligands exhibited stabilizing effects on the enzyme, particularly in regions critical to ligand binding, such as the active site residues. Residues in proximity to the active site, including LEU 264, ASN 130, and HIS 128, showed reduced fluctuations in the ligand-bound forms compared to the unbound enzyme, suggesting tighter interactions and less conformational freedom [17]. This aligns with the role of these residues in forming key interactions with the ligands during docking.

The radius of gyration (Rg) analysis (Figure 3.3a) revealed consistent structural compactness of the enzyme when bound to VS-1 and VS-2, compared to the unbound form. A more compact structure, as indicated by lower Radius of gyration values, correlates with greater structural stability, and both ligand-bound forms maintained compactness throughout the simulation [18]. This suggests that the ligands did not induce significant structural deviations in the enzyme, supporting the possibility that the interactions with VS-1 and VS-2 contribute to maintaining the enzyme’s integrity during the simulation.

The number of hydrogen bond interactions between the ligands and the enzyme was a critical factor in stabilizing the *T.b* TR-ligand complexes. Over the 100 ns simulation time, both VS-1 and VS-2 exhibited stable hydrogen bond interactions with key residues in the enzyme’s active site (Figure 3.4). VS-2 formed an average of 1–3 hydrogen bonds throughout the simulation, primarily with residues such as HIS 128 and ASN 130. These bonds were critical for maintaining the ligand’s position within the active site and contributed to the overall stability of the complex [19]. VS-1, on the other hand, formed slightly more hydrogen bonds (2–3 on average), and the interactions were stronger, particularly with residues ASN 261, MET 260, and VAL 133, as predicted from docking studies. These hydrogen bonds played a pivotal role in anchoring VS-2 within the active site, explaining its higher binding free energy compared to VS-1 [19]. The consistent hydrogen bond formation throughout the simulation suggests that both ligands form strong, stable interactions with the enzyme, supporting their potential as effective inhibitors.

The solvent accessible surface area (SASA) analysis provided additional insights into the exposure of the enzyme and ligand surfaces to the solvent, which is critical for understanding the solvation effects and the potential for ligand binding [20]. Over the course of the simulation, both ligand-bound forms of *T.b* TR exhibited lower SASA values compared to the unbound enzyme (Figure 3.5), indicating that ligand binding reduced the solvent exposure of the enzyme’s active site. For the *T.b* TR-VS1 complex, the SASA value consistently decreased as the simulation progressed, reflecting the stable and buried nature of the ligand within the enzyme’s binding pocket [20]. This reduced solvent exposure suggests that VS-1 occupies a large portion of the active site, shielding key residues from solvent interactions and stabilizing the complex.

Similarly, in the *T.b* TR-VS2 complex, a significant reduction in SASA was observed, although the decrease was slightly less pronounced than in the VS-1 complex. This indicates that VS-2 also fits well within the enzyme’s active site but leaves some regions more exposed to the solvent compared to VS-1 [20]. Nonetheless, the lower SASA values for both complexes indicate that the ligands effectively bind to and protect the enzyme’s active site from solvent interactions, contributing to the overall stability of the complexes.

### Binding Free Energy Calculation

To further evaluate the strength of the interactions between *T.b* TR and the ligands, binding free energy calculations were performed using the MMPBSA (Molecular Mechanics Poisson-Boltzmann Surface Area) approach [21]. The binding free energy values of the *T.b* TR-VS1 and *T.b* TR-VS2 complexes were −30.93 kcal/Mol and −32.93 kcal/Mol, respectively (Figures 3.6a and 3.7a). These values indicate favorable binding affinities, with VS-2 exhibiting slightly stronger binding compared to VS-1. Both ligands demonstrated higher binding free energies than Melarsoprol (the standard drug), which suggests that they may be more effective inhibitors of *T.b* TR [22]. The binding free energy primarily arises from electrostatic and van der Waals interactions, which were significant contributors to the overall binding affinity. This is consistent with the docking results, where both ligands formed stable hydrogen bonds and hydrophobic interactions with key residues in the enzyme’s active site. The solvation energy (ΔGsolv) also contributed favorably to the binding free energy, since ligand-enzyme complexes are well-solvated in the aqueous environment used during simulations.

Per-residue decomposition of the binding free energy provided insights into the contributions of individual amino acid residues to the overall stability of the protein-ligand complexes [23]. In the *T.b* TR-VS1 complex (Figure 3.6b), MET 260, ASN 130 and ILE 110 contributed the most to the binding energy, with MET 260 contributing the highest (−5.09 kcal/Mol). These residues, located in the enzyme’s active site, were crucial for stabilizing VS-1 through hydrophobic and van der Waals interactions [23]. The high contribution of MET 260 indicates that it plays a pivotal role in the binding of VS-1, reinforcing its importance in the enzyme-ligand interaction. For the *T.b* TR-VS2 complex (Figure 3.7b), ASN 130 and LEU 129 were the most significant contributors, with ASN 130 contributing −2.01 kcal/Mol to the binding free energy. These residues form critical hydrogen bonds with VS-2, as identified during the docking analysis, and their large contributions to the binding energy highlight the importance of electrostatic interactions in stabilizing the complex. The strong binding affinity of VS-2 can be attributed to these specific interactions, which anchor the ligand firmly within the enzyme’s active site.

## Conclusion

In this study, the pharmacophore model derived from Melarsoprol was successfully validated and used for structure-based virtual screening to identify potential inhibitors of *Trypanosome brucei* Trypanothione Reductase (*T.b* TR). The top two compounds, VS-1 and VS-2, exhibited strong binding affinities, as indicated by their superior docking scores compared to Melarsoprol. Molecular dynamics simulations further confirmed the stability of both complexes, with favorable hydrogen bonding and solvent accessible surface area (SASA) profiles. Per-residue decomposition identified key amino acids that contribute significantly to ligand binding, providing valuable insights into the interactions required for effective inhibition of *T.b* TR. Overall, these results suggest that both VS-1 and VS-2 are promising drug candidates for the treatment of Human African Trypanosomiasis (HAT), showing higher potential than the current standard treatment, Melarsoprol.

## Limitations and Recommendations

Carrying out molecular dynamics simulations of the melarsoprol complexed with *Trypanosome brucei* Trypanothione Reductase enzyme was not possible. During melarsoprol topology generation step using melarsoprol mol2 file, the arsenic atom (As) in Melarsoprol was not parametizable in CHARMM General Force Field (CGenFF) to generate a string file which is needed in the ligand topology generation. We recommend conducting *in vitro* assays to validate the in silico findings for VS-1 and VS-2. This might include the determination of the IC50 values of both compounds against *Trypanosome brucei gambiense* and *Trypanosome brucei rhodesiense* to assess their potency. Additionally, this can be supplemented with a cell cycle analysis using flow cytometry to check for cell cycle arrest and a morphological assay to observe structural changes in the parasites post treatment. To ensure selectivity, determination of the selectivity index can be carried out by comparing the compounds’ cytotoxicity against mammalian cells versus the parasites. Lastly, performance a growth kinetics assay to monitor the parasites’ growth in the presence of the compounds over time. These experiments will confirm the efficacy and safety of VS-1 and VS-2 for potential use against Human African Trypanosomiasis.

## Supporting information

All the python scripts used in this work are accessible via (https://github.com/Elcanah402/drug_discovery.git).

## Acknowledgements

The authors gratefully acknowledge the use of the ZUPUTO High Performance Computing facility, hosted by the West Africa Center for Cell Biology of Infectious Pathogens (WACCBIP), for providing computational resources. The molecular dynamics simulations in this study were performed on 7212 X NVIDIA TESLA P100 16GB GPUs, and we sincerely appreciate the support and infrastructure made available for this work.

## Author Contributions

**Conceptualization:** Elcanah Mauta Evans

**Methodology**: Elcanah Mauta Evans, Oluwaseun oluwatosin Taofeek, Theresa Manful Gwira,

**Data Analysis:** Elcanah Mauta Evans, Collins Misita Morang’a, Thommas Mutemi Musyoka, Steven Ger Nyanjom

**Writing**: Elcanah Mauta Evans, Oluwaseun oluwatosin Taofeek

